# AVF creation in mice disrupts blood flow patterns to impact vasoreactivity

**DOI:** 10.64898/2026.11.11.766047

**Authors:** Sohom Mookherjee, Hannah Northrup, Revi Brown, Nathan Hill, Bing Li, Timmy Lee, Shelley Wang, Sihem Boudina, Yan-Ting Shiu, J. David Symons

**Affiliations:** Department of Nutrition and Integrative Physiology and Molecular Medicine Program, University of Utah, Salt Lake City, UT; Veterans Affairs Medical Center, Salt Lake City, UT; Division of Nephrology and Hypertension, Department of Internal Medicine, University of Utah, Salt Lake City, UT; Nora Eccles Harrison Cardiovascular Research and Training Institute, University of Utah; Department of Medicine and Division of Nephrology, University of Alabama at Birmingham, AL; Veterans Affairs Medical Center, Birmingham, AL

**Author notes:** Address for correspondence: Sohom Mookherjee.

**Keywords:** endothelium, vascular smooth muscle, dialysis, chronic kidney disease, arterio-venous fistula

## Abstract

Maladaptive venous remodeling contributes to the high rate of arteriovenous fistula (AVF) maturation failure. Impaired vasoreactivity might impact this pathology. We created a murine AVF by anastomosing the external jugular vein to the carotid artery, establishing the AVF vein (AVF-V) and proximal (PA) and distal (DA) carotid artery. We hypothesized that endothelium-dependent vasorelaxation is impaired in the AVF-V vs. the contralateral vein (contra-V) of 5-month-old mice of both sexes 21-days post-surgery, reasoning that abrupt exposure to arterial pressure and disturbed hemodynamics promote endothelial dysfunction. Further, because magnetic resonance imaging revealed lower blood flow velocity and wall shear stress in PA vs. DA 21-days post-surgery, we hypothesized that endothelial function is compromised in PA. Whereas the thromboxane A2 receptor (TP) agonist U-46619 evoked sufficient vasocontraction in the contra-V from which subsequent vasorelaxation could be evaluated, AVF-V responsiveness was insufficient for this assessment, potentially reflecting >5-fold increases in collagen content and intima-media thickness. Consistent with our arterial hypothesis, endothelium-dependent vasorelaxation was impaired in the PA vs. DA and contralateral artery, whereas vascular smooth muscle function, cross-sectional area, collagen area, and intima-media thickness were similar among segments. Low shear stress impairs endothelial autophagy, which supports endothelial nitric oxide generation and vasodilation. Colocalized fluorescence of p-eNOS^S1177^ with CD31+ cells, and autophagy-related protein 3 with CD31+ cells, as well as nitric oxide bioavailability, were lower in PA vs. DA. These findings suggest that differential blood flow patterns in PA vs. DA following AVF creation may modulate endothelial autophagy, nitric oxide generation, and endothelium-dependent vasorelaxation.

**Graphical abstract:** 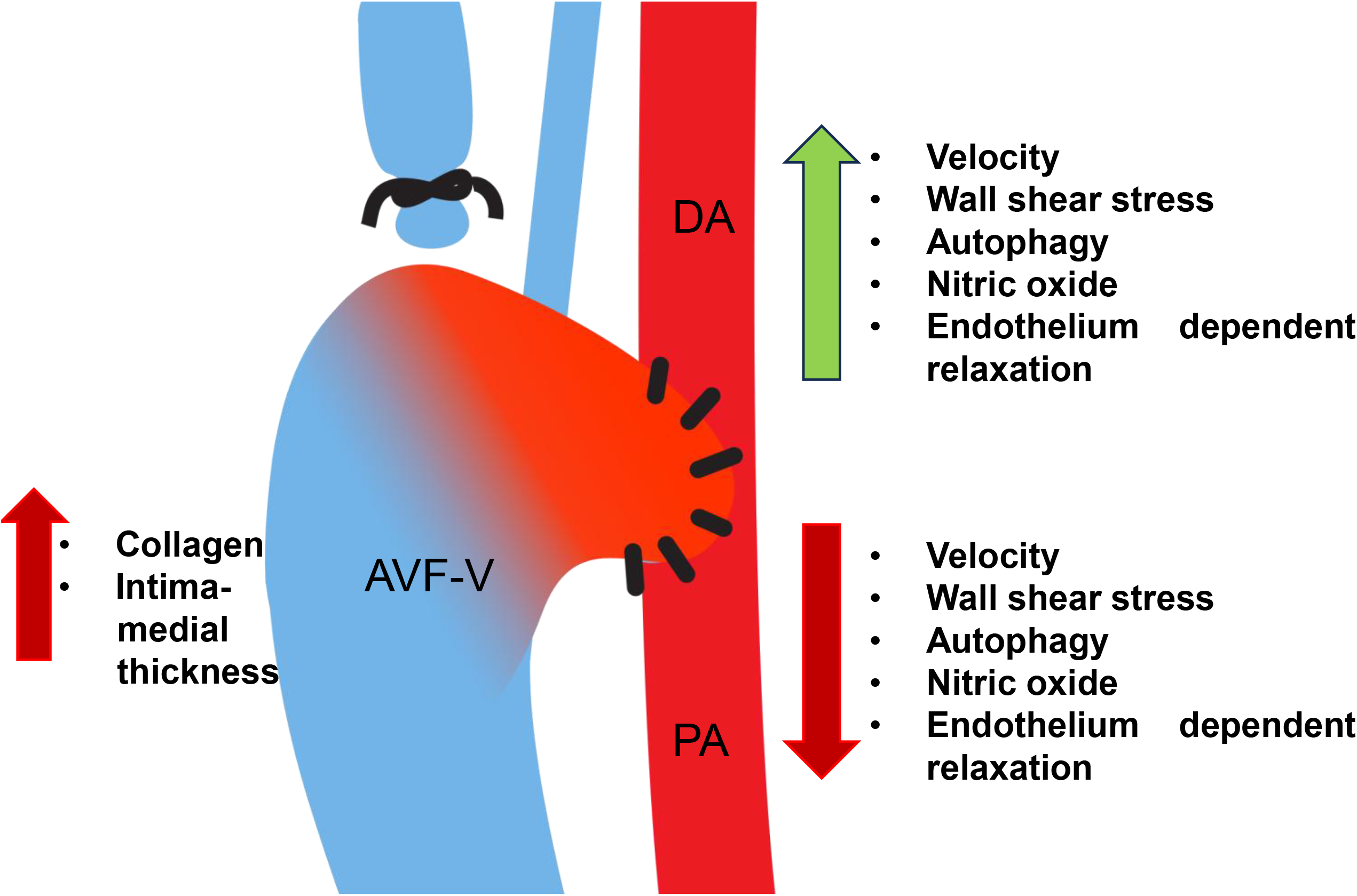

## Introduction

The arteriovenous fistula (AVF) is the preferred vascular access for hemodialysis in patients with end-stage kidney disease.(1) Despite this preference, up to 60% of AVFs created in the United States fail to mature adequately for dialysis use.(2) AVF maturation failure is attributed largely to maladaptive venous remodeling, characterized by excessive intimal hyperplasia and inadequate lumen expansion; however, the mechanisms underlying these processes remain incompletely understood.(3)

One working hypothesis is that endothelial and/or vascular smooth muscle dysfunction within the venous and arterial components of the AVF could initiate or amplify inappropriate vascular remodeling and thereby contribute to AVF failure.(3) We employ a commonly used murine model wherein the external jugular vein is anastomosed to the carotid artery to establish three AVF components: the AVF vein (AVF-V) and the arterial segments proximal (PA) and distal (DA) to the anastomosis. Using approaches previously established by our laboratory to assess endothelial and vascular smooth muscle function in each component,(4) we tested the hypotheses that endothelium-dependent vasorelaxation is impaired in the AVF-V relative to the contralateral vein (contra-V), and in the PA vs. DA and contralateral artery (contra-A).

Limited U-46619–induced vasocontraction in the AVF-V relative to the contra-V 21-days post AVF surgery precluded reliable assessment of endothelium-dependent vasorelaxation and, consequently, testing our first hypothesis. The inability to respond likely reflected marked collagen accrual and intima-media thickening observed in the AVF-V vs. contra-V. Magnetic resonance imaging (MRI) of vessel segments at 21-days post AVF creation revealed lower blood flow velocity and arterial wall shear stress in PA vs. DA and contra-A. Based on these findings, we hypothesized endothelium-dependent vasorelaxation would be compromised in PA vs. DA and contra-A and results were supportive. Preliminary and published (5–8), (9) reports from our laboratory and others indicate reduced flow velocity and shear stress compromise endothelial autophagy in arterial and venous segments, and that intact autophagy contributes importantly to nitric oxide (NO) production by arterial endothelial cells. Immunohistochemistry revealed that colocalized fluorescence of p-eNOS^S1177^ with CD31+ cells and autophagy-related protein 3 (ATG3) with CD31+ cells, as well as NO bioavailability, were lower in the PA vs. DA, whereas vascular smooth muscle function was similar between segments. Differential blood flow patterns in PA vs. DA following AVF creation could modulate AVF arterial function, at least in part, through an endothelial autophagy–eNOS–NO signaling pathway. A preliminary report of these findings has been presented. (10)

## Methods

### Ethical approval

All procedures were approved by the University of Utah Institutional Animal Care and Use Committee (IACUC), the Salt Lake City Veterans Affairs Medical Center IACUC, and the University of Alabama at Birmingham IACUC. Methodologies were conducted in accordance with the National Institutes of Health *Guide for the Care and Use of Laboratory Animals*, and complied with established ethical guidelines.(11)

### Animals and housing

Male and female C57BL/6J mice were purchased from The Jackson Laboratory (Bar Harbor, ME) at 3 months of age and housed four per cage in the AAALAC-accredited Veterinary Medical Unit (VMU) at the Salt Lake City Veterans Affairs Medical Center, which is operated under contract by the Comparative Medicine Center at the University of Utah, or the Animal Resources Program (ARP) at the University of Alabama at Birmingham (22–23°C; 12-h light/12-h dark cycle). Mice received standard rodent chow (Envigo, Catalog #2920X, Indianapolis, IN) and reverse-osmosis, autoclaved water *ad libitum* and were monitored daily.

### Carotid-jugular AVF creation

Anesthesia was induced with 2–5% isoflurane in 100% oxygen and maintained with 1–3% isoflurane in 5-month-old male (n=3) and female (n=4) mice. Mice were positioned supine on a heating pad, and an end-to-side AVF was created by anastomosing the external jugular vein to the carotid artery.(12, 13) Experiments were performed 21-days after AVF creation. Mice used for vascular function, histology, and immunofluorescence experiments were housed at the VMU, whereas mice used for MRI studies were housed at the ARP.

### Magnetic resonance imaging (MRI)

Upon a stable plane of anesthesia, MRI was performed in 3-month-old male C57BL/6J mice.(12, 14) Images were acquired using two-dimensional T2-weighted fast spin-echo with black-blood double inversion for lumen and two-dimensional gradient-echo velocity mapping for velocity.

Three-dimensional AVF lumen reconstructions were generated from black-blood images using Amira 3D 2021.1 (Thermo Fisher Scientific, Waltham, MA). Following smoothing, flow extensions were added at vessel boundaries to minimize entrance effects. Meshing was performed using ANSYS ICEM CFD 2020 R2 (Ansys, Inc., Canonsburg, PA) and consisted of 1–1.5 million tetrahedral elements with four prism layers at the vessel wall.

Blood flow velocity was determined using ImageJ. PA and AVF-V velocities were prescribed as velocity boundary conditions, and the DA was assigned as a pressure outlet with 0 gauge pressure. CFD simulations were performed using ANSYS Fluent 2020 R2 with rigid, no-slip vessel walls. Blood was modeled as Newtonian and incompressible, with a density of 1050 kg/m³ and viscosity of 0.0035 Pa·s. Residual convergence was set at 1 × 10 . Simulations were run for three cardiac cycles, with the final cycle used for analysis.

Post-processing was performed using Tecplot 360 EX 2019 R1 (Tecplot, Inc., Bellevue, WA). Vessel centerlines were calculated using VMTK at 0.1-mm increments, and centerline-normal coordinates were calculated in MATLAB (MathWorks, Natick, MA). These coordinates were used to determine velocity magnitude (Eq. 1), wall shear stress (WSS; Eq. 2), and average cross-sectional area of the PA and AVF-V from 0–40 mm from the anastomosis. Values were compared with those of the DA at the MRI measurement site, ∼20 mm from the anastomosis. Data were analyzed using GraphPad Prism (GraphPad Software, San Diego, CA).

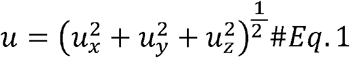

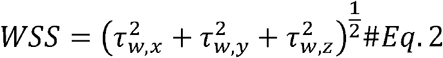

### Vascular responsiveness

Mice were anesthetized as described above, the chest was opened, and 1 mL physiological saline solution (PSS; pH 7.35–7.40) was perfused through the left ventricle after incision of the right atrium. The AVF-V, PA, DA, contra-V, and contra-A were isolated in ice-cold PSS and mounted in a wire myograph using two 20-µm tungsten wires, one secured to a stationary micrometer and the other to a force transducer. Chambers were heated from room temperature to 37°C over 45-min with vessels maintained at 0 mg tension.

For arteries, a length-tension relationship was established to determine Lmax, followed by concentration-response curves to U-46619 (10□□–10□□ M). Arteries were then precontracted with U-46619 to ∼65% of maximal tension, and endothelium-dependent vasorelaxation to acetylcholine (ACh; 10□□–10□□ M) was assessed in the absence and presence of the nitric oxide (NO) synthase inhibitor N^G^-monomethyl-L-arginine acetate (L-NMMA; 10□³ M). Endothelium-independent vasorelaxation to sodium nitroprusside (SNP; 10□□–10 □□M) was assessed similarly. Between concentration-response curves, vessels were washed immediately and at 10-min intervals for 30 min. For veins, a modified length-tension relationship was used to determine Lmax, followed by U-46619-induced vasocontraction and assessment of vasorelaxation as described above and in detail elsewhere.(4)

### Data acquisition

Vascular tension was recorded continuously using an analog-to-digital interface (Biopac Systems Inc., Santa Barbara, CA) for offline analysis. These procedures have been used to assess arterial and venous reactivity in clinical(15) and preclinical(8) studies.

### Histology

The AVF-V, contra-V, PA, DA, and contra-A were formalin-fixed, paraffin-embedded, and sectioned at 5 µm. Sections were stained with Masson’s trichrome (MT) and Verhoeff–Van Gieson (VVG) to assess collagen deposition and vascular morphology, respectively. With MT, smooth muscle stains red and collagen blue; collagen deposition was calculated as blue-stained medial area/total medial area × 100%. With VVG, elastic fibers and lamellae stain black, permitting visualization of the internal and external elastic laminae and delineation of vascular layers. Sections were imaged by bright-field microscopy and analyzed using Bioquant software.(16)

### Immunofluorescence

Arterial and venous sections were deparaffinized by heating at 60°C for 30 min, incubation in CitriSolv (Decon Labs, Cat#1601), and graded ethanol washes (100%, 95%, 70%, and 50%) followed by ddH O. Antigen retrieval was performed using 1% Antigen Unmasking Solution (Vector Laboratories, Cat#H-3301). Sections were blocked for 1 h in 2.5% donkey serum/PBS, incubated with primary antibodies overnight at 4°C and fluorophore-conjugated secondary antibodies for 2 h at room temperature, followed by DAPI (Invitrogen, Cat#62248) for 5-min and mounting (Invitrogen, Cat#P36934). Antibodies and dilutions are listed in Supplemental Table 2. Images were acquired using a Leica SP8 confocal microscope.(6–8, 17)

Immunofluorescence was quantified using manual annotation and CellProfiler.(18) ECs were delineated using CD31, and mean fluorescence intensity of ATG3 and p-eNOS^S1177^ colocalized with CD31+ ECs was quantified as previously described.(6–8, 17)

### Statistical analyses

Values are expressed as means ± SEM. Single endpoints were compared between: two vessel segments (i.e., AVF-V vs. contra-V) via paired t-test; or among three vessel segments (i.e., PA, DA, contra-A) via one-way ANOVA, followed by Tukey’s post hoc test when the overall ANOVA was significant (*P*<0.05). Concentration-response data were compared using repeated-measures two-way ANOVA, followed by Tukey’s post hoc tests when significant interaction effects were detected.

## RESULTS

### MRI

At 21-days post AVF surgery, blood flow velocity (m•s^-1^) and wall shear stress (dyne•cm^2^) were lower in the PA and AVF-V vs. DA **(Figure 1 A-F)**. The AVF-V cross sectional area was markedly larger vs. both arterial components of the AVF **(Figure 1G)**.

**Figure 1.**
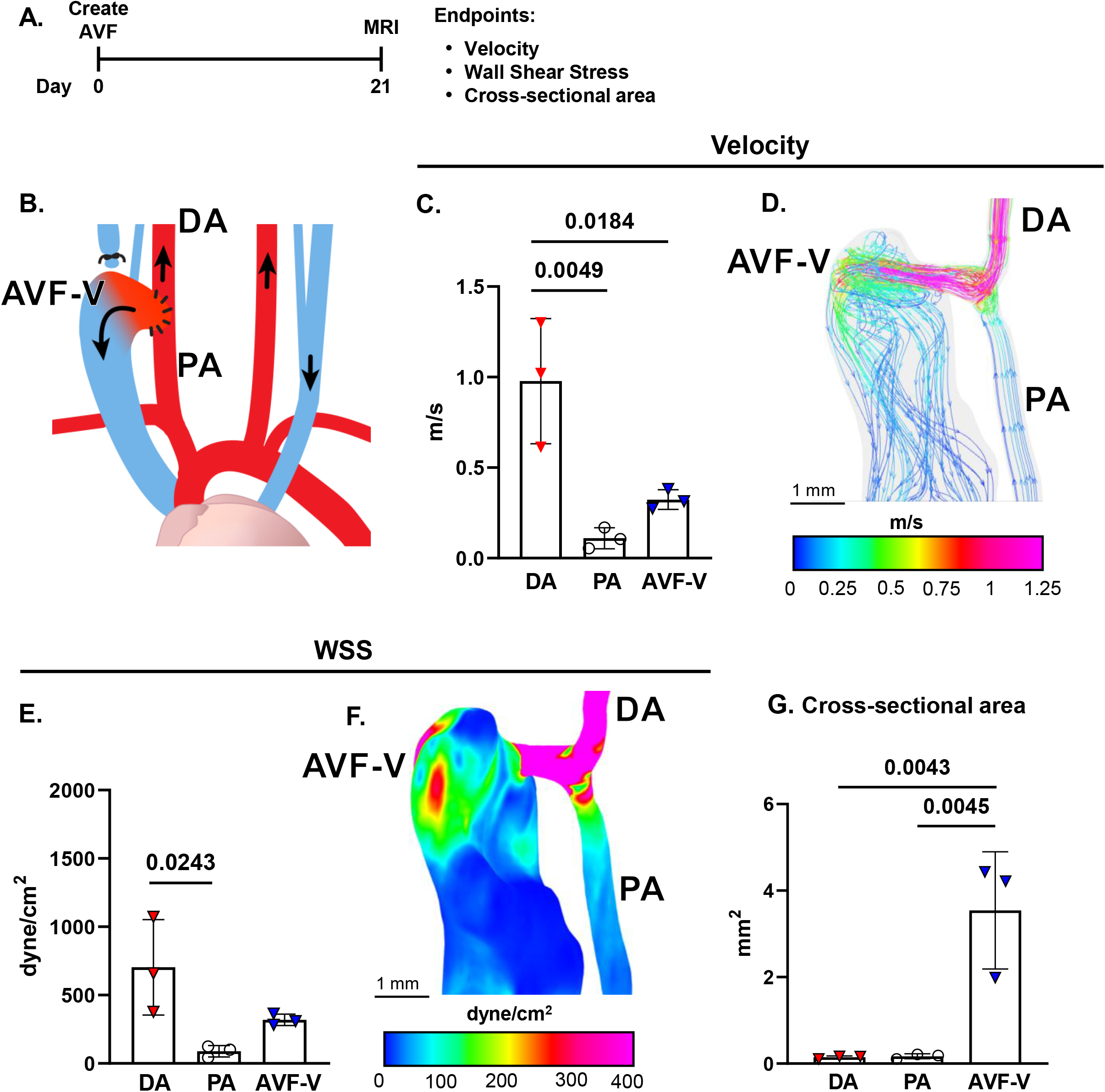
Magnetic resonance imaging (MRI). **A.** MRI was completed 21-d post AVF creation in the **(B)** AVF-vein (AVF-V), distal artery (DA) and proximal artery (PA). From these measures, mean values **(C)** and representative images **(D)** of blood flow velocity, together with mean values **(E)** and representative images **(F)** of wall shear-stress (WSS) were obtained. MRI also enabled cross-sectional area of each AVF component to be calculated **(G)**. Data represent mean ± SE for 3 mice. Comparisons were made among groups via one-way ANOVA. When a significant p value was obtained (p<0.05), the location of the difference was identified using a Tukey’s post hoc test.

### Vascular responsiveness and histology – AVF-V and contra-V

Start-width, end-width, and vessel length were not different between the AVF-V and the contra-V **(Supplemental Table 1)**. U-46619 is a stable synthetic analog of the endoperoxide prostaglandin PGH_2_ and was used as a TP agonist. U-46619-mediated TP activation couples with and mobilizes one or more G proteins to activate phospholipase C, IP3, and intracellular Ca^2+^ in vascular smooth muscle cells. Relative to baseline tension, U-46619-evoked tension development (mg tension / µm length) occurred in both venous segments, but maximal responses were markedly greater in the contra-V (0.036±0.006 mg tension / µm length) vs. the AVF-V (0.016±0.006 mg tension / µm length). Due to limited contraction displayed by the AVF-V, the assessment of endothelium-dependent vasorelaxation could not be made in a reliable manner. Blunted vasocontraction might reflect our observation that collagen area and intima-media thickness were markedly higher in the AVF-V vs. the contra-V **(Figure 2A-F)**. In the contra-V, maximal Ach-induced vasorelaxation was 43±6% and maximal SNP-induced vasorelaxation was 55±5%, suggesting strongly that the inability to reliably assess vasorelaxation in the AVF-V was due to a biological rather than technical limitation.

**Figure 2.**
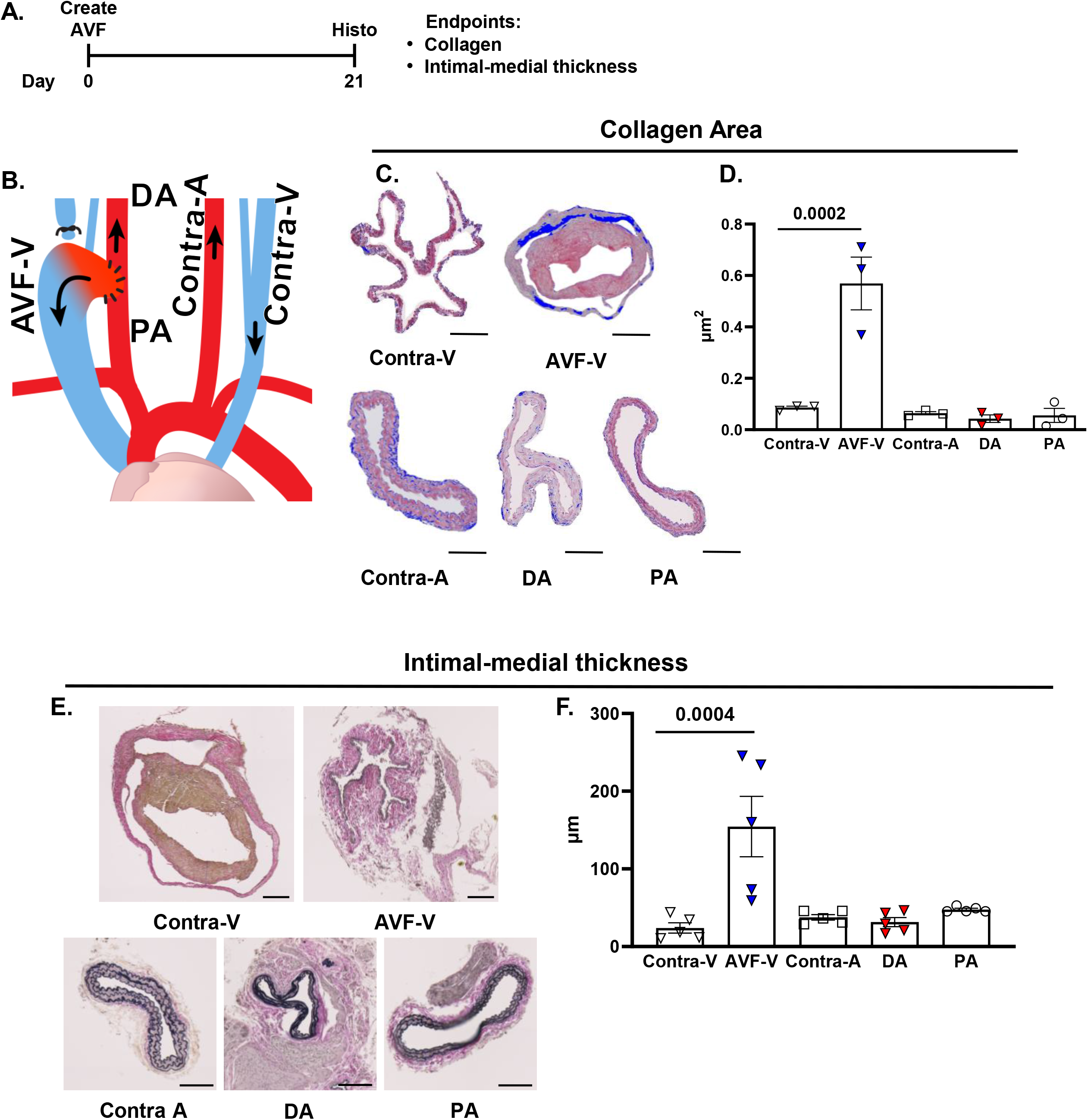
Histology. **A.** Histology was completed 21-d post AVF creation in the **(B)** AVF-vein (AVF-V), distal artery (DA) and proximal artery (PA), together with the contralateral artery (contra-A) and vein (contra-V). Representative images **(C, E)** of mean data **(D, F)** for collagen area and intimal-medial thickness are shown. Data represent mean ± SE for 3-5 mice. Comparisons were made among groups via one-way ANOVA. When a significant p value was obtained (p<0.05), the location of the difference was identified using a Tukey’s post hoc test. Scale bar: 100 um.

#### Vascular responsiveness and histology – PA, DA, and contra-A

Start-width, end-width, and vessel length were not different among the PA, DA, or contra-A **(Supplemental Table 1)**. Relative to baseline tension, U-46619-evoked vasocontraction was similar between the PA and DA **(Figure 3A-C)**, but lower vs. the contra-A **(Supplemental Figure 1A-C)**.

**Figure 3.**
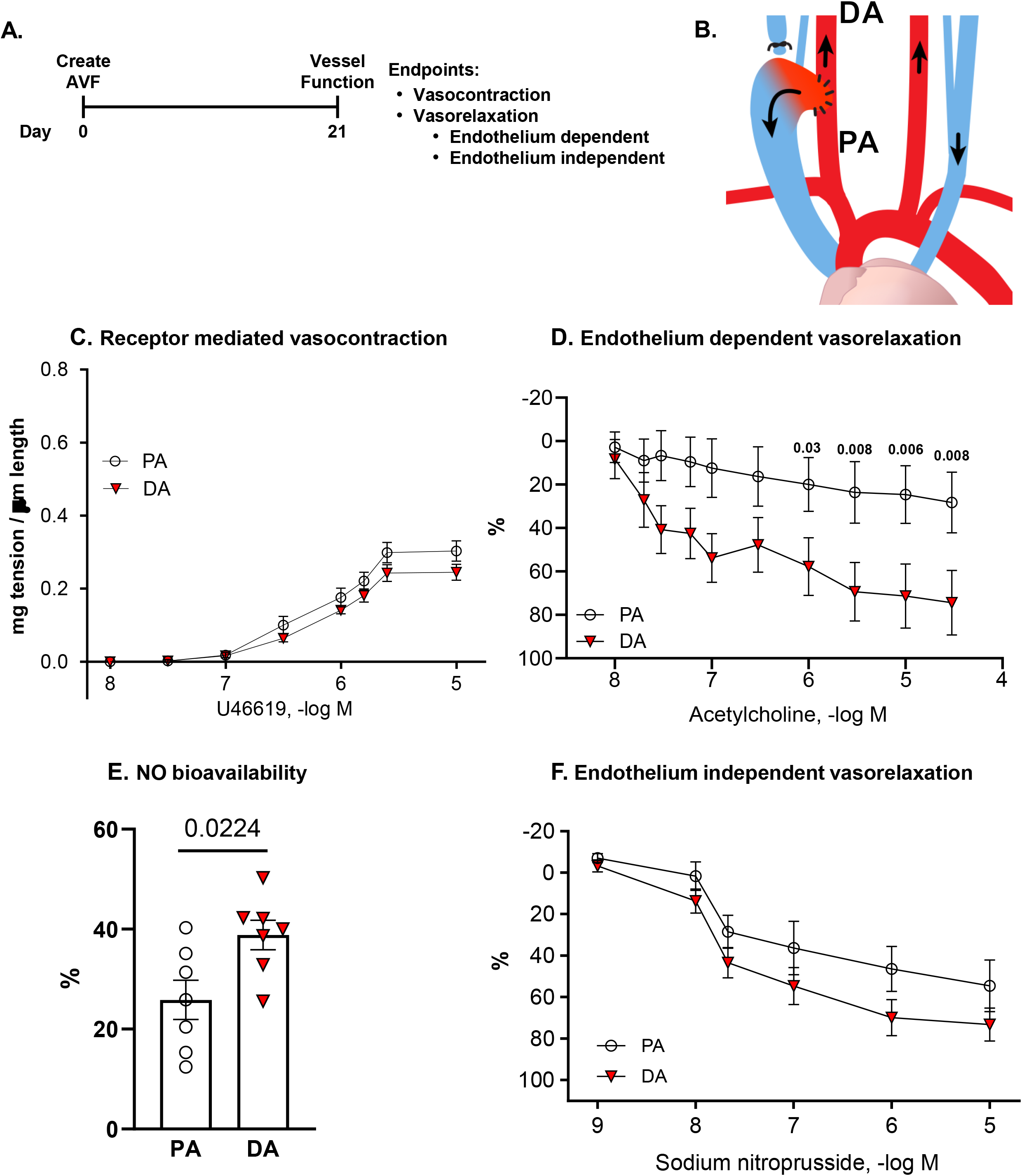
Vascular responses. **A.** Vascular reactivity was assessed 21-d post AVF creation in the **(B)** arterial segments proximal (PA) and distal (DA) to the anastomosis. **C.** Vasocontraction to the thromboxane A_2_ receptor agonist U-46619, **(D)** vasorelaxation to acetylcholine (Ach), **(E)** nitric oxide (NO) bioavailability, and **(F)** vasorelaxation to sodium nitroprusside (SNP) are shown. Comparisons were made among groups via two-way repeated measures ANOVA for C, D, F. When a significant p value was obtained (p<0.05), the location of the difference was identified using a Tukey’s post hoc test. Results shown in Panel E were compared via unpaired t-test. Data represent mean ± SE from 7 mice x 1 vessel segment per mouse.

ACh binds to M_3_ muscarinic receptors on the vascular endothelium to increase intracellular Ca^2+^. Elevated intracellular Ca^2+^ then activates constitutive type III NOS, which facilitates conversion of the amino acid substrate L-arginine to the products L-citrulline and NO. In arterial segments stably precontracted using U-46619, ACh evoked vasorelaxation was lower in the PA vs. DA **(Figure 3D)**. Maximal responses to Ach in DA were similar to those observed in contra-A **(Supplemental Figure 1D)**.

To assess the contribution from NO to vasorelaxation in the PA, DA, and contra-A, we completed ACh concentration – response curves after vessels incubated for 30-min with L-NMMA. L-NMMA competes with the endogenous NOS substrate L-arginine. Because L-NMMA is administered to the vessel bathing medium in excess, it outcompetes L-arginine for NOS binding sites and thereby limits the capability for NO generation. Maximal ACh-evoked vasorelaxation was attenuated in the presence (i.e., +L-NMMA) vs. the absence (i.e., -L-NMMA) of NOS inhibition in the DA and contra-A but not the PA **(Supplemental Figure 2A-E)**. These results were used to calculate NO bioavailability.(19) As would be predicted based on these responses and Ach findings displayed in **Figure 3D and Supplemental Figure 1D**, NO bioavailability was limited in the PA vs. the DA and contra-A **(Figure 3E, Supplemental Figure 2F).**

NO that is produced by endothelial cells diffuses abluminally to the vascular smooth muscle where it activates guanylyl cyclase (GC). GC stimulation increases cGMP formation which inhibits Ca^2+^ entry into the vascular smooth muscle and precipitates vasorelaxation. SNP is a nitrodilator that releases NO spontaneously to activate GC directly. As such, SNP is used to assess endothelium-independent vasorelaxation. After precontraction using U-46619, SNP-evoked vasorelaxation was similar in PA and DA **(Figure 3F)** and did not differ when compared to results obtained in the contra-A **(Supplemental Figure 3A-C)**. In contrast to the AVF-V and contra-V, no differences existed among the PA, DA, and contra-A regarding collagen area or intima-media thickness **(Figure 2E-F)**.

### Immunofluorescence

Based on preliminary results and published reports from our laboratory group and others,(1, 12, 13, 16, 20, 21) we tested the hypothesis that vascular dysfunction observed in the AVF-V vs. the contra-V associates with lower p-eNOS^S1177^ and autophagy related protein 3 (ATG3) expression. In partial support, ATG3 colocalization with CD31+ cells (i.e., ECs) trended lower (p=NS) whereas p-eNOS^S1177^ colocalization with CD31+ cells was decreased in AVF-V vs. contra-V **(Supplemental Figure 4).**

Regarding the arterial components of the AVF, data reported by others (22), (23) and ourselves (5–8, 24) motivated the hypothesis that lower blood flow velocity, wall shear stress, endothelium-dependent vasorelaxation, and NO bioavailability observed in the PA vs. DA associates with reduced ATG3 expression and p-eNOS^Ser1177^. Consistent with this, immunohistochemistry results indicate ATG3 and p-eNOS^Ser1177^ colocalization with CD31+ cells was lower in the PA vs. DA **(Figure 4A–F)**.

**Figure 4.**
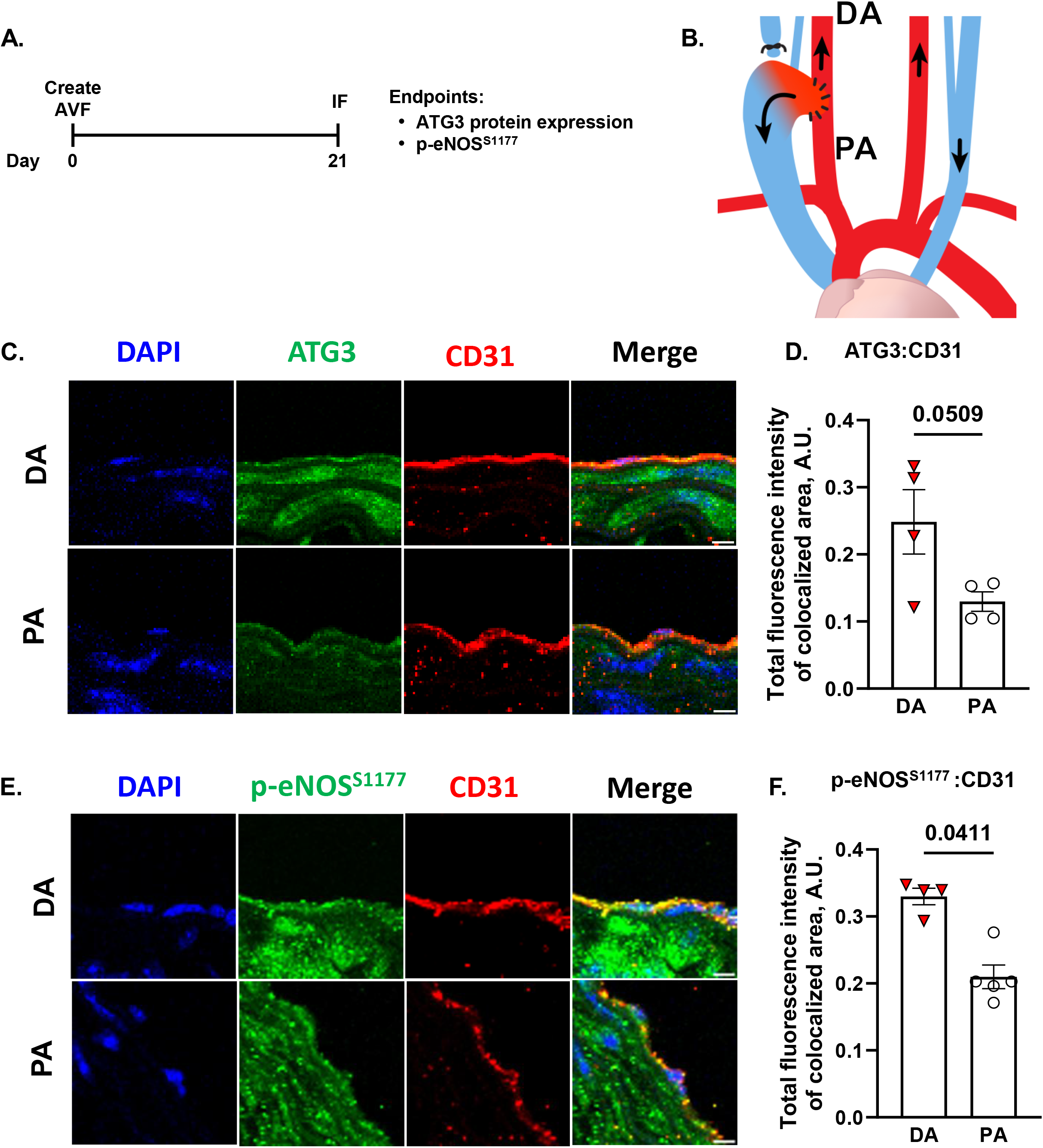
Immunofluorescence (IF). **A.** IF was completed 21-d post AVF creation in the **(B)** arterial segments proximal (PA) and distal (DA) to the anastomosis. Images **(C, E)** of mean data **(D, F)** for total fluorescence intensity of ATG3 or p-eNOS^S1177^ colocalized with CD31+ cells is shown. For C, DAPI (blue), antibodies to autophagy-related gene 3 (ATG3; green), and antibodies to endothelial cell surface marker CD31 (red) were used to visualize nuclei, ATG3, and endothelial cells, respectively. For E, all stains were similar to C except that antibodies to activating phosphorylation of eNOS at serine 1177 (p-eNOS^S1177^; green) were used instead of ATG3. Comparisons were made via unpaired t-test. Significance was obtained when p<0.05. Data in D and F represent mean ± SE from 4-5 mice (limited by tissue avalibility). Each point in the histogram represents the mean for a single mouse obtained from 3 sections per mouse and 3 fields of view per section. All CD31+ cell in each field of view were analyzed. Magnification = 63x. Scale bar = 5 um.

## DISCUSSION

Understanding the pathobiology of AVF maturation failure is essential for developing effective therapies to reduce morbidity and mortality in patients requiring hemodialysis vascular access. This pathology is commonly characterized by stenosis near the venous anastomosis, excessive intimal hyperplasia, and inadequate outward remodeling.(3) Endothelial and/or vascular smooth muscle dysfunction within the venous and arterial components of the AVF may contribute to these adverse outcomes. Here we report three principal findings. First, although endothelial function could be assessed in the contralateral vein (contra-V), insufficient precontraction of the AVF vein (AVF-V) precluded reliable assessment of endothelium-dependent vasorelaxation. Second, because MRI revealed lower blood flow velocity and wall shear stress in the proximal (PA) vs. distal (DA) arterial segment, we hypothesized that endothelium-dependent vasorelaxation would be impaired in the PA relative to the DA; our findings supported this hypothesis. Third, vascular dysfunction in the AVF-V and PA was associated with reduced endothelial indices of autophagy and p-eNOS^S1177^. Collectively, these findings suggest that regional differences in AVF hemodynamics may modulate endothelial autophagy, eNOS signaling, and vascular function.

### AVF-V vs. contra-V

In an earlier study we characterized endothelial and vascular smooth muscle function of the murine external jugular vein using isometric myography.(4) Here we used this approach to test the hypothesis that endothelial function is impaired in the AVF-V vs. contra-V. This hypothesis was based on the abrupt exposure of the normally low-pressure venous circulation to an altered hemodynamic environment characterized by heightened blood flow and changes in intraluminal pressure, circumferential wall stress, and spatially heterogeneous wall shear stress.(21, 25) These mechanical forces can disrupt endothelial homeostasis and contribute to vascular dysfunction and structural adaptation of the AVF vein.(20) Further rationale was provided by our review of the literature, which identified no previous assessment of vasoreactivity in the murine AVF-V.

Maximal U-46619-induced vasocontraction and ACh-induced vasorelaxation in the contra-V (0.040 mg tension/µm length and 43 ± 6%, respectively) were comparable to values we previously reported in external jugular veins of naïve mice (0.036 mg tension/µm length and 58 ± 7%, respectively).(4) Here, in contrast, the AVF-V exhibited minimal U-46619-induced vasocontraction, precluding reliable assessment of endothelium-dependent vasorelaxation. The concordance between responses of the contra-V in the present study and external jugular vein in our earlier investigation(4) supports the interpretation that the attenuated AVF-V contractile response reflects vascular dysfunction rather than a technical limitation.

A potential contributor to impaired U-46619-induced vasocontraction is the extensive structural remodeling of the AVF-V. AVF creation increased collagen content and intima-media thickness approximately fivefold, consistent with extracellular matrix accumulation and vascular wall remodeling. Martinez et al. demonstrated that postoperative venous medial fibrosis is associated with AVF maturation failure,(26) whereas Zhao et al. showed that arterialization induces medial thickening and vascular smooth muscle cell phenotypic modulation within the neointima.(27) Thus, extracellular matrix accumulation and wall thickening could alter vessel mechanics and reduce the translation of TP receptor-mediated vascular smooth muscle cell contraction into measurable force, whereas smooth muscle cell phenotypic modulation could compromise the contractile apparatus itself. Importantly, our experiments do not establish a causal relationship between collagen accumulation and impaired TP receptor-mediated contraction. Accordingly, structural remodeling may contribute to the attenuated U-46619 response, while altered TP receptor signaling, smooth muscle cell phenotypic modulation, impaired Ca²□ handling or actomyosin force generation, and increased mechanical constraint imposed by extracellular matrix accumulation remain plausible, non-mutually exclusive mechanisms.

We initially hypothesized that endothelium-dependent vasorelaxation would be impaired in the AVF-V vs. contra-V based on preliminary and published observations from our laboratory and others. In this regard, AVF creation exposes venous endothelium to altered and spatially disturbed hemodynamic forces.(12, 13, 21) Consistent with potential consequences of this environment, Franzoni et al. reported reduced p-eNOS^S1177^ in human umbilical vein endothelial cells (HUVECs) exposed to disturbed vs. laminar shear stress.(20) In addition, previous studies from our laboratory (5–8) and others (9, 22, 23) indicate that flow and shear stress regulate endothelial autophagy and that intact autophagy supports endothelial NO bioavailability. Together, these observations motivated us to determine whether endothelial ATG3 and p-eNOS^S1177^ are reduced in the AVF-V vs. contra-V. Although inadequate AVF-V precontraction prevented reliable assessment of endothelium-dependent vasorelaxation, immunofluorescence findings were partially supportive: ATG3 colocalization with CD31+ cells trended lower, whereas p-eNOS^S1177^ colocalization with CD31+ cells was reduced, in the AVF-V vs. contra-V.

### PA, DA, and contra-A

Using MRI procedures described previously,(12, 13) we observed lower blood flow velocity and arterial wall shear stress in the PA vs. DA 21 days after AVF creation. These findings inspired us to test if endothelium-dependent vasorelaxation would be impaired in the PA relative to the DA. Consistent with this hypothesis, ACh-induced vasorelaxation and NO bioavailability were attenuated in the PA vs. DA, whereas SNP-induced vasorelaxation was similar between segments, supporting an endothelial rather than vascular smooth muscle defect. Moreover, endothelium-dependent and -independent vasorelaxation were similar between the DA and contra-A, indicating that endothelial function was impaired in the PA but preserved in the DA.

Differential shear-dependent regulation of endothelial autophagy and eNOS/NO signaling provides a plausible mechanism for the divergent PA and DA responses. Laminar shear promotes endothelial autophagy and eNOS expression, phosphorylation, and NO generation, whereas low or disturbed shear is associated with impaired autophagic activity and endothelial dysfunction (5, 9, 22, 23, 28). Thus, the higher blood flow velocity and wall shear stress in the DA may favor maintenance of endothelial autophagy and eNOS/NO signaling, consistent with preserved ACh-induced vasorelaxation and NO bioavailability. Conversely, the lower-shear environment of the PA may provide less stimulus for these protective pathways, contributing to impaired endothelium-dependent vasorelaxation. Preservation of SNP-induced vasorelaxation further supports an endothelial defect rather than impaired vascular smooth muscle responsiveness to NO. We therefore determined whether the lower flow velocity, shear stress, and endothelial function in the PA versus DA were accompanied by reduced endothelial autophagy and eNOS activation. Immunohistochemical analysis demonstrated reduced colocalization of ATG3 and p-eNOS^S1177^ with CD31+ cells in the PA relative to the DA. Collectively, these findings support the concept that regional differences in arterial hemodynamics after AVF creation are associated with corresponding differences in endothelial function, potentially through autophagy–eNOS–NO signaling.

#### Experimental considerations

Several points should be considered when integrating our findings into the current literature. First, although we studied adult mice of both sexes, sample size precluded the evaluation of sex-dependent differences. This is relevant because evidence exists that AVFs mature less frequently and more slowly in women than in men,(29, 30) and that sex-dependent differences in arterial inflow and vascular remodeling might contribute to these divergent outcomes. (31) Second, AVFs were created in otherwise naïve mice which therefore lacked the metabolic milieu associated with chronic kidney disease (CKD). This is important because reports exist that the CKD milieu exacerbates neointimal hyperplasia formation in AVFs, likely, at least in part, through CKD-induced endothelial dysfunction. (32–35) The impact of manipulating whole-body or EC-selective eNOS or ATG3 expression has yet to be determined in mice in the absence and presence of CKD. Third, whether and how CKD modifies eNOS and autophagic responses to laminar vs. disturbed flow in venous and arterial ECs remains unknown. In conclusion, our report provides proof-of-concept evidence supporting more comprehensive investigation of endothelial autophagy as a potential contributor to AVF maturation. Ongoing studies by our laboratory group are evaluating the three considerations mentioned above and others, together with specific nodes of the autophagic pathway and direct measures of autophagic flux to further explore the potential for endothelial autophagy to mitigate AVF maturation failure.

## Acknowledgements

The authors thank Yuxia He and Tatyana Isayeva Waldrop for assisting animal work performed in Utah and Alabama, respectively.

## Data Availability

The raw data supporting the conclusions of this manuscript will be made available by the authors, without undue reservation, to any qualified researcher.

## Funding Disclosure

Support was provided for: SM by University of Utah (UU) Graduate Research Fellowship and American Heart Association (AHA) Predoctoral Fellowship AHAPRE1025910; RB by the Medical Student Research Program at the UU Spencer Fox Eccles School of Medicine, supported by the Eccles Endowment for Medical School Excellence; NH by R25HL161608; TL, YTS, and JDS by MPI R01HL153244; SB by 1R01HL184520-01-A1 and 1R01HL167866-01A1; YTS by R01DK129299, VA I01BX006269, VA I01BX006078, and Nora Eccles Treadwell Foundation Project 51004733; and JDS by 1RO1HL141540.

## Ethics Statement

The Institutional Animal Care and Use Committee of the University of Utah approved these experiments.

## Author Contributions

SM, JDS, YTS, SB, and TL designed the study. SM, HN, RB, NH, BL, SW, and JDS performed experiments. SM and JDS wrote the manuscript. All authors contributed to editing and revising.

## Conflict of Interest Statement

TL is a consultant for Proteon Therapeutics, Merck, and Boston Scientific. The other co-authors declare no competing interests.

## Supplemental figure legends

**Supplemental Figure 1.**
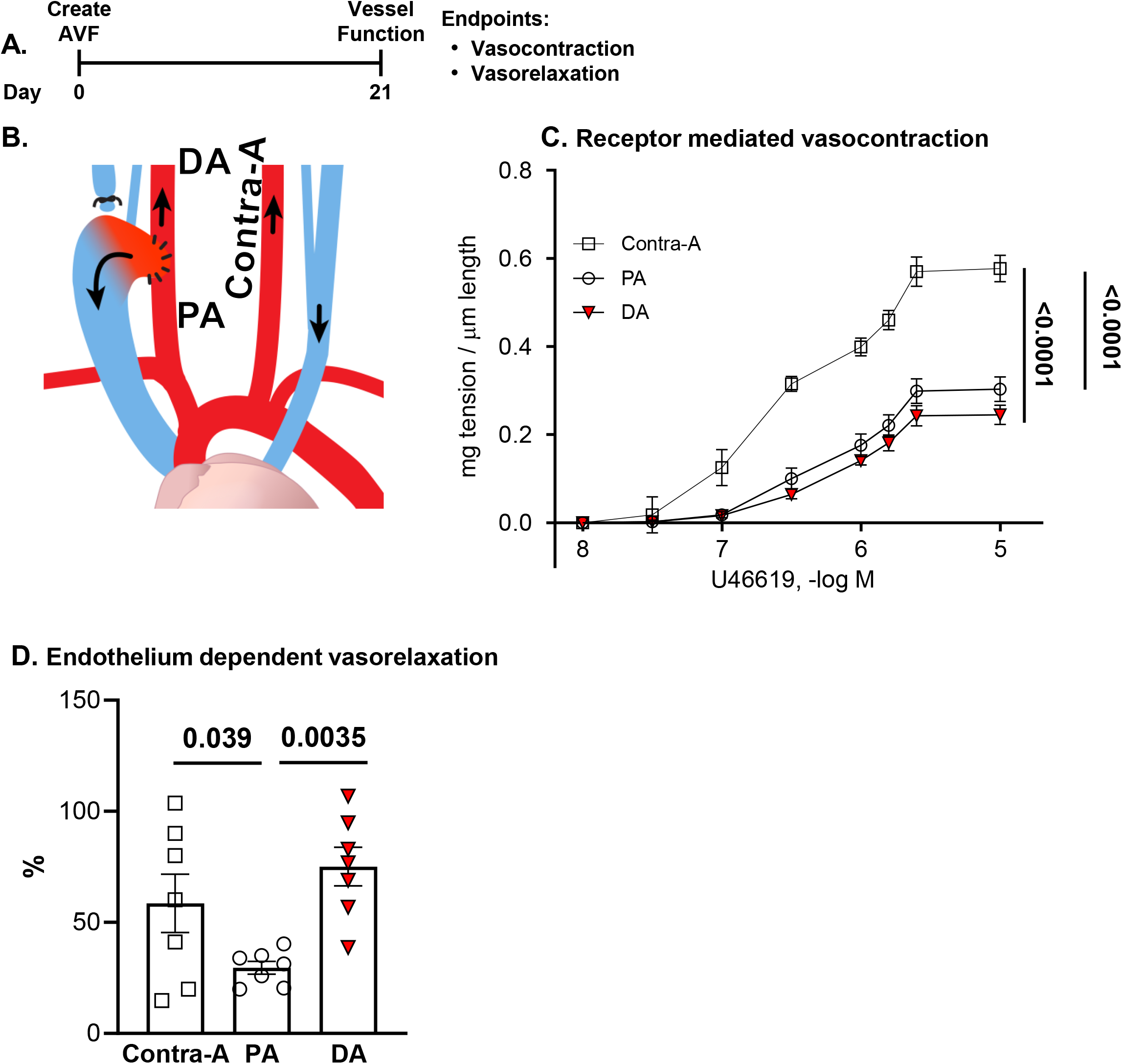
Vascular responses. **A.** Vascular reactivity was assessed 21-d post AVF creation in the **(B)** arterial segments proximal (PA) and distal (DA) to the anastomosis together with the contralateral artery (contra-A). **C.** Vasocontraction to the thromboxane A_2_ receptor agonist U-46619 and **(D)** maximal vasorelaxation to acetylcholine (ACh) are shown. Comparisons were made among groups via two-way repeated measures ANOVA for C and one-way repeated measures ANOVA for D. When a significant p value was obtained (p<0.05), the location of the difference was identified using a Tukey’s post hoc test. Data represent mean ± SE from 7 mice x 1 vessel segment per mouse.

**Supplemental Figure 2.**
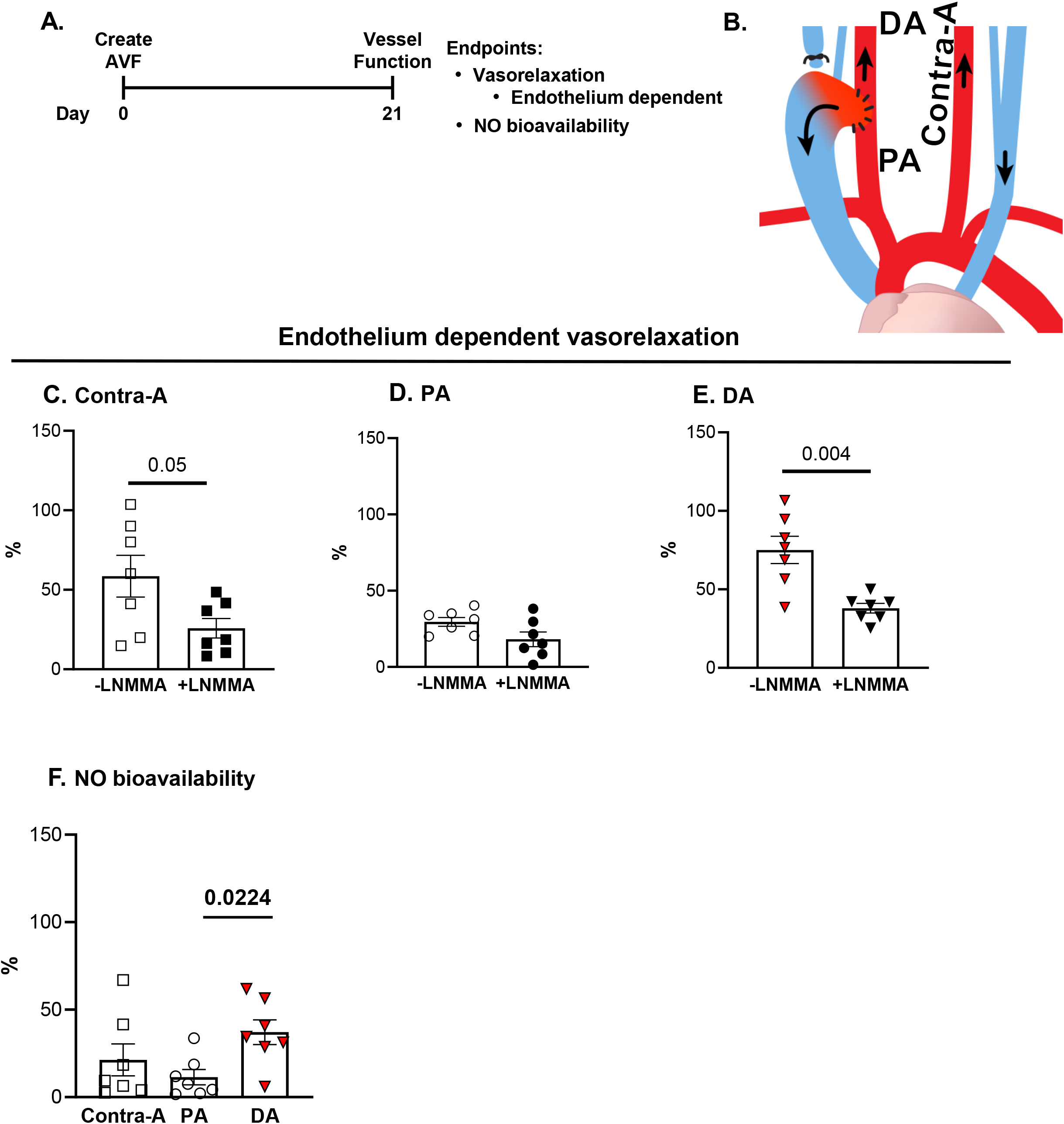
Nitric oxide bioavailability. **A**. Endothelium dependent vasorelaxation was assessed 21-d post AVF creation in the **(B)** arterial segments proximal (PA) and distal (DA) to the anastomosis together with the contralateral artery (contra-A). **C-F:** Maximal vasorelaxation to acetylcholine (Ach) in the presence and absence of LNMMA in the contra-A **(C)**, PA **(D)**, and DA **(E)** were used to calculate NO bioavailability **(F)**. Comparisons were made between conditions using an unpaired t-test in panels C – E. Results shown in panel F were obtained via one-way repeated measures ANOVA. A Tukey’s post-hoc test was used to identify the location of difference. Data represent mean ± SE from 7 mice x 1 vessel segment per mouse.

**Supplemental Figure 3.**
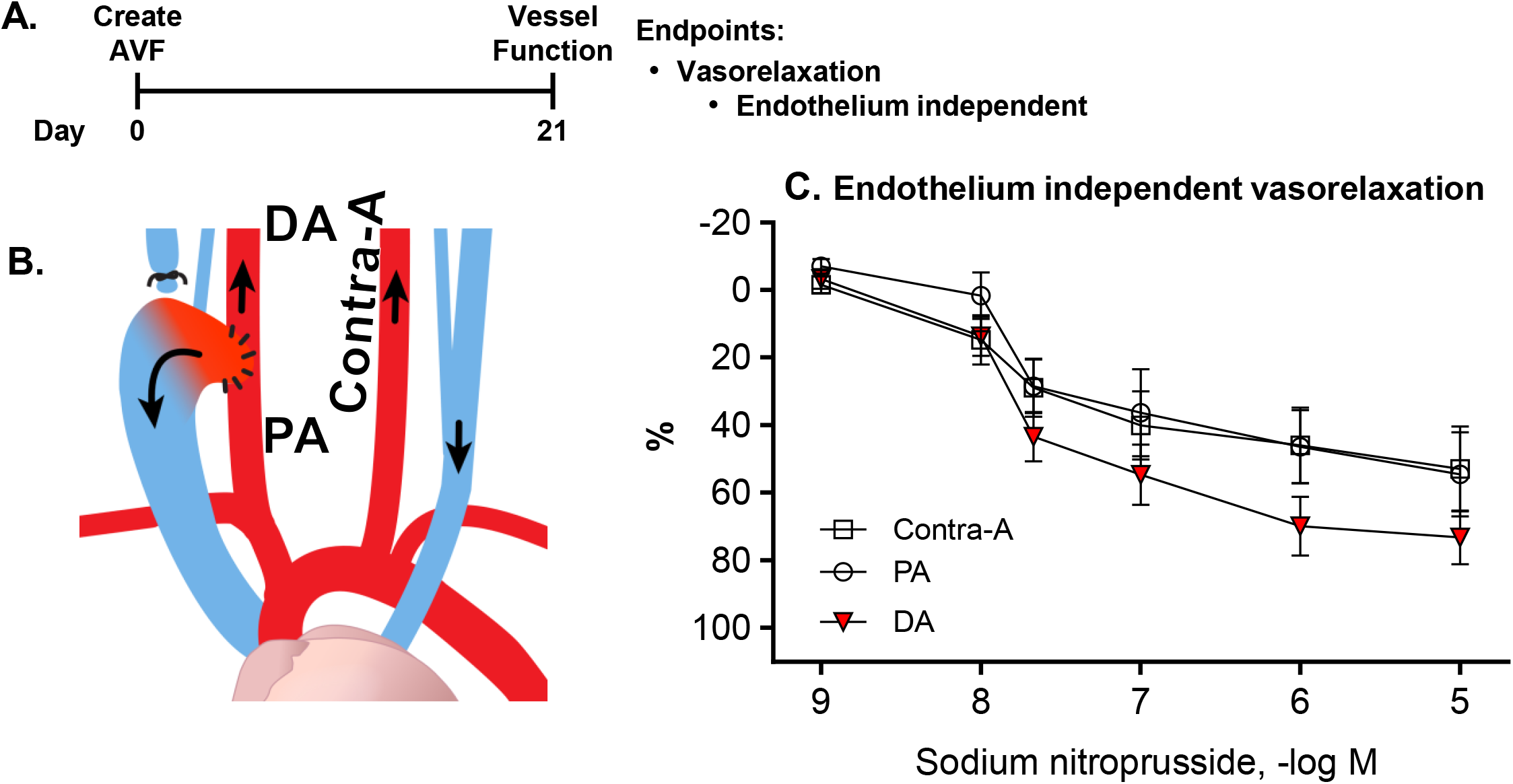
Smooth muscle responses. **A.** Endothelium independent vasorelaxation was assessed 21-d post AVF creation in the **(B)** arterial segments proximal (PA) and distal (DA) to the anastomosis together with the contralateral artery (contra-A). **C.** Vasorelaxation to sodium nitroprusside (SNP). Comparisons were made among groups via two-way repeated measures ANOVA. Data represent mean ± SE from 7 mice x 1 vessel segment per mouse.

**Supplemental Figure 4.**
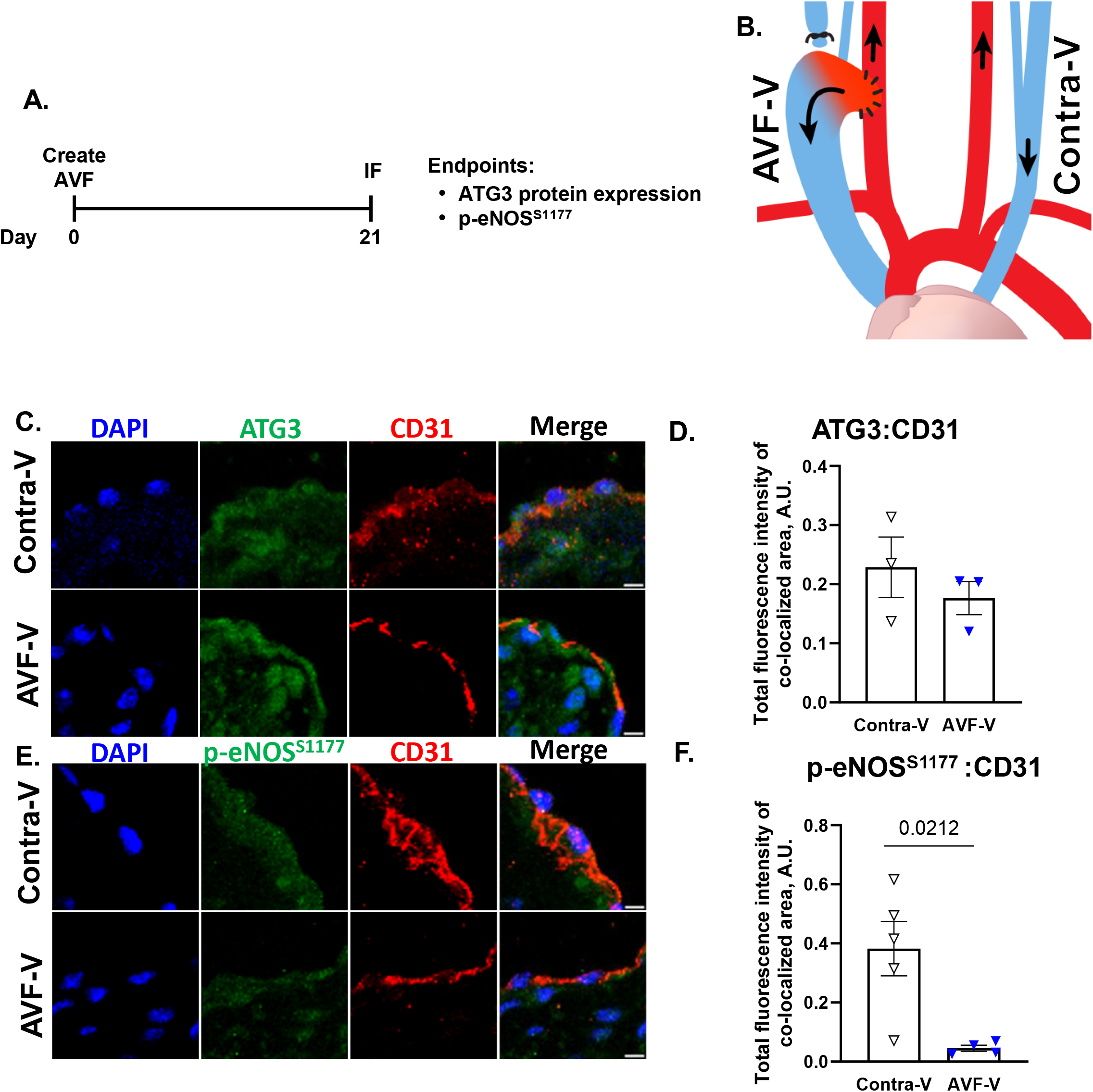
Immunofluorescence (IF). **A.** IF was completed 21-d post AVF creation in the **(B)** venous segment of the arteriovenous fistula (AVF-V) and the contralateral side (contra-V). Images **(C, E)** and mean data **(D, F)** for total fluorescence intensity of ATG3 or p-eNOS^S1177^ colocalized with CD31+ cells is shown. For C, DAPI (blue), antibodies to autophagy-related gene 3 (ATG3; green), and antibodies to endothelial cell surface marker CD31 (red) were used to visualize nuclei, ATG3, and endothelial cells, respectively. For E, all stains were similar to C except that antibodies to activating phosphorylation of eNOS at serine 1177 (p-eNOS^S1177^; green) were used instead of ATG3. Comparisons were made via unpaired t-test. Significance was obtained when p<0.05. Data in D and F represent mean ± SE from 3 mice per group (limited by tissue avalibility). Each point in the histogram represents the mean for a single mouse obtained from 3 sections per mouse and 3 fields of view per section. All CD31+ cell in each field of view were analyzed. Magnification = 63x. Scale bar = 5 um.

**Supplemental Table 1:** Vessel Characteristics. Start width, end width, and length in um of vessel segments from AVF vein (AVF-V), contralateral vein (contra-V), proximal artery (PA), distal artery (DA) and contralateral artery (contra-A). Data represent mean ± SE from 7 mice x 1 vessel segment per mouse.

**Supplemental Table 2:** Source, catalog number, and host by target protein for antibodies used in immunofluorescence staining and the dilution that was used.

## Notes

### Competing Interest Statement

The authors have declared no competing interest.

